# sureLDA: A Multi-Disease Automated Phenotyping Method for the Electronic Health Record

**DOI:** 10.1101/2020.04.13.038968

**Authors:** Yuri Ahuja, Doudou Zhou, Zeling He, Jiehuan Sun, Victor M. Castro, Vivian Gainer, Shawn N. Murphy, Chuan Hong, Tianxi Cai

## Abstract

**Objective:** A major bottleneck hindering utilization of electronic health record (EHR) data for translational research is the lack of precise phenotype labels. Chart review as well as rule-based and supervised phenotyping approaches require laborious expert input, hampering applicability to studies that require many phenotypes to be defined and labeled *de novo*. Though ICD codes are often used as surrogates for true labels in this setting, these sometimes suffer from poor specificity. We propose a fully automated topic modeling algorithm to simultaneously annotate multiple phenotypes.

**Methods:** sureLDA is a label-free multidimensional phenotyping method. It first uses the PheNorm algorithm to initialize probabilities based on two surrogate features for each target phenotype, and then leverages these probabilities to constrain the Latent Dirichlet Allocation (LDA) topic model to generate phenotype-specific topics. Finally, it combines phenotype-feature counts with surrogates via clustering ensemble to yield final phenotype probabilities.

**Results:** sureLDA achieves reliably high accuracy and precision across a range of simulated and real-world phenotypes. Its performance is robust to phenotype prevalence and relative informativeness of surogate versus non-surrogate features. It also exhibits powerful feature selection properties.

**Discussion:** sureLDA combines attractive properties of PheNorm and LDA to achieve high accuracy and precision robust to diverse phenotype characteristics. It offers particular improvement for phenotypes insufficiently captured by a few surrogate features. Moreover, sureLDA’s feature selection ability enables it to handle high feature dimensions and produce interpretable computational phenotypes.

**Conclusion:** sureLDA is well suited toward large-scale EHR phenotyping for highly multi-phenotype applications such as PheWAS.

## INTRODUCTION

Electronic health records (EHR), often linked with biorepositories, have become an increasingly important data source for translational research. [1, 2] Such rich, multidimensional data promise myriad opportunities for translational applications ranging from personalizing care decisions to predicting disease prognosis. [3] However, the scarcity of precise phenotype labels has hampered efforts to harness this dataset for many of these objectives. Studies focusing on one or a few phenotypes typically circumvent this problem by predicting labels from diverse EHR features using 1) rule-based algorithms or 2) supervised learning methods trained on a subset of manually anontated gold-standard labels (GLabels). [4, 5, 6, 7, 8, 9, 10, 11] Of note, the Phenotype Knowledgebase (PheKB) platform built for the Electronic Medical Records and Genomics (eMERGE) Network has been shown to effectively integrate expertise across sites to yield accurate, transportable phenotyping algorithms. [11] However, applying these approaches to new phenotypes requires substantial expert input: manually annotating GLabels to train supervised methods necessitates laborious chart review, and formulating rule-based algorithms involves iteratively devising and validating rules. Thus these approaches, while effective, are infeasible for highly multi-phenotype applications such as Phenome-Wide Association Studies (PheWAS) requiring *de novo* labeling of hundreds to thousands of phenotypes. This signifies an ongoing need for phenotyping methods that can simultaneously annotate many diverse phenotypes.

Currently, studies requiring many EHR phenotypes often utilize billing codes such as International Classification of Diseases (ICD) codes as surrogates for true phenotype labels. For instance, the original PheWAS [12, 13, 14] grouped ICD codes into ∼1800 phenotype codes (phecodes) that were subsequently thresholded and screened for associations with various genetic markers. While a trivial function of ICD codes can be reliably used in lieu of true labels for many diseases, others such as rheumatoid arthritis [15] tend to have inprecise codes that can diminish the power of the association study. [16] Thresholding ICD codes at higher counts boosts label PPV and specificity but may significantly diminish sensitivity, especially for episodic conditions such as pseudogout.

To produce more accurate phenotype labels without extensive expert intervention, researchers have recently advocated for label-free (i.e. not requring GLabels) computational phenotyping methods. [17, 18, 19, 20, 21, 22, 23, 24] The class of “weakly supervised” methods, which train supervised classifiers using noisy labels generated from key surrogate features in the data rather than expensive GLabels, has proven particularly powerful to this end. For instance, the “anchor and learn” approach trains a regularized logistic regression model on imperfect labels derived from ‘anchor’ features with high PPVs. [17, 18] While this approach avoids GLabels, it requires expert input to identify appropriate anchors. Several other efforts have fully automated the pipeline by using knowledge-base derived labels or standardized “silver standard” surrogates, such as the ICD code or NLP mention for the target phenotype, to train classifiers. For instance, Levine et al. demonstrate high concordance between lagged linear regression models trained using a knowledge-base derived standard versus an expert clinician derived one. [24] Similarly, the XPress method predicts a phenotype by fitting a regularized logistic regression on a noisy label defined as the presence of at least one key ICD code. [19] The PheNorm method uses multiple surrogates derived from the main ICD and NLP by 1) fitting log-normal mixtures adjusting for healthcare utilization, and 2) performing additional denoising with other features via dropout training. [20] Likewise, the MAP algorithm fits a multimodal clustering ensemble to the ICD and NLP surrogates adjusting for healthcare utilization. [21]

These automated methods often achieve impressive accuracy and can be practically scaled to a large number of phenotypes. However, we hypothesize two means of improvement: 1) better incorporating information from non-surrogate features, and 2) jointly predicting phenotypes in order to impose an Occam’s Razor [25] assumption that fewer concurrent diseases is more likely than more. To achieve these goals we draw inspiration from the UPhenome algorithm - a modification of the widely-used Latent Dirichlet Allocation (LDA) topic model [26] that combines diverse features to infer patients’ joint distributions over unbiased phenotype ‘topics’ [27] Though UPhenome was developed for phenotype discovery rather than annotation of known phenotypes, the “grounded phenome” (GPhenome) algorithm derived therefrom better reflects known phenotypes by setting topic priors to the pseudocount of clinical terms associated with each phenotype, thus portending potential application to an annotation task. [28]

In this paper we propose surrogate-guided ensemble LDA (sureLDA), an automated multi-phenotype annotation method that employs probabilistic pseudolabels produced by PheNorm to guide topic inception in LDA in a “weakly supervised” manner. As with GPhenome this guiding of LDA is inspired by two well-trodden LDA derivatives, labeled LDA and semi-supervised LDA, [29, 30] though using only surrogates rather than GLabels to constrain topics. Unlike GPhenome, sureLDA 1) utilizes phenotype probabilities from PheNorm rather than clinical concept counts as “noisy” labels to guide topic formation in LDA, 2) weights features using regression coefficients from the dropout training step of PheNorm, and 3) employs a cluster ensemble approach to combine guided LDA scores with surrogates to yield posterior phenotype probabilities. sureLDA combines desirable properties of PheNorm and LDA, effectively leveraging the typically informative surrogate features using PheNorm while exploiting LDA’s prowess at extracting information from high-dimensional feature spaces to jointly annotate multiple phenotypes. Consequently, we submit that sureLDA is uniquely well-suited to high-throughput phenotyping of the EHR for a highly multi-phenotype application akin to PheWAS.

## METHODS

The sureLDA algorithm broadly consists of four key steps: (i) assemble informative features, including the main ICD and NLP features as silver-standard surrogates for each target phenotype, as well as a healthcare utilization feature (*H*); (ii) use PheNorm to obtain initial probabilities for each target phenotype based on these surrogates together with *H*; (iii) fit guided LDA to all features simultaneously using the initial probabilities as Dirichlet hyperparameters for the patient-phenotype distributions, yielding patient-level phenotype “scores”; and (iv) perform ensemble clustering of the surrogates and the LDA phenotype scores (again adjusting for *H*) to predict final probabilities for each target phenotype. A schematic illustrating this procedure is displayed in Figure 1. Throughout, we assume that there are a total of *K* target phenotypes with true binary status denoted by **Y** = (*Y*_1_,. .., *Y*_*K*_)’, which again is not used to train sureLDA. Let **X** denote the entire feature vector of dimension *p* and **X**^*log*^ = log(**X** + 1). We assume there are a total of *D* patients in the EHR study and use subscript *d* to index the patients.

**Figure 1:**
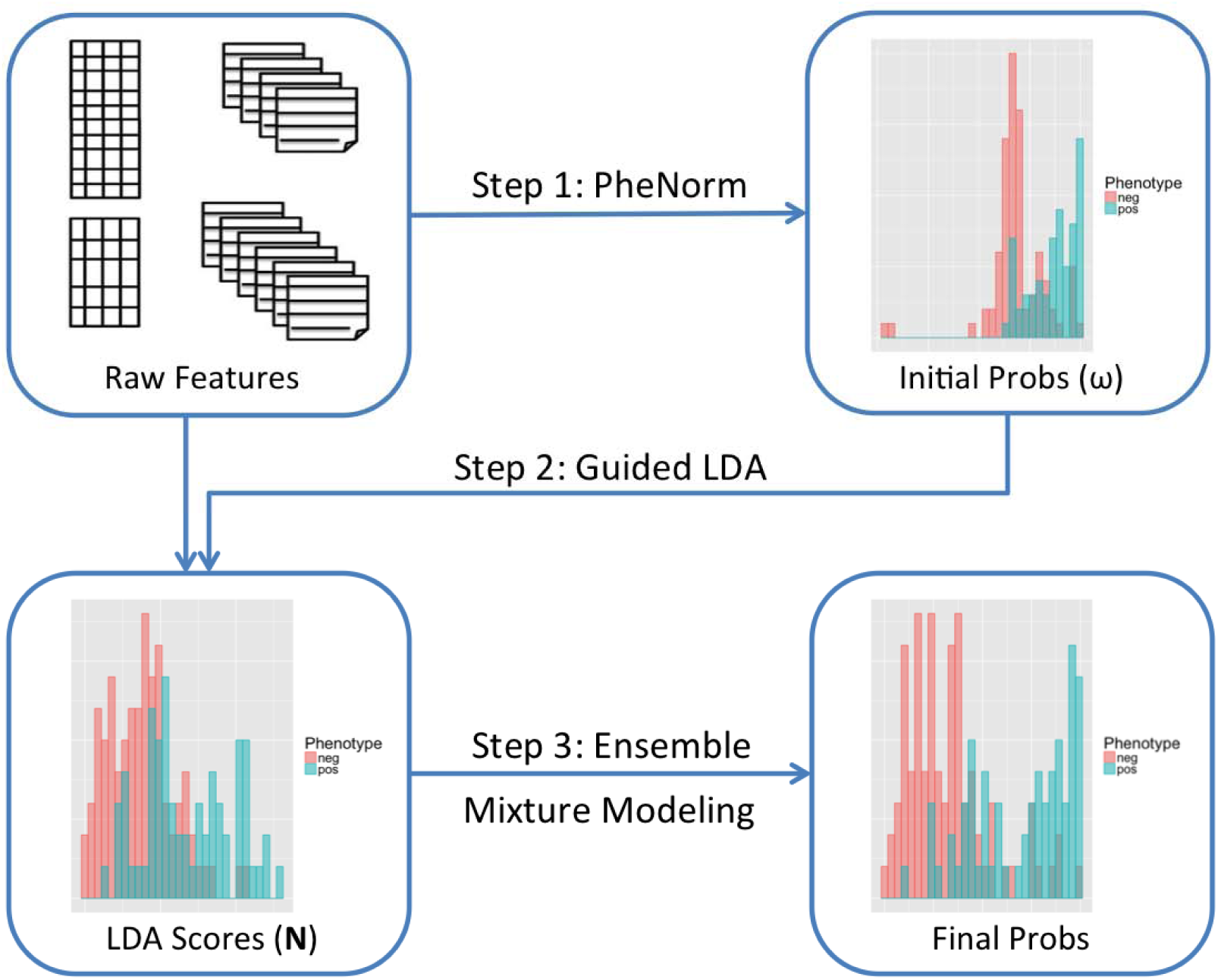
Schematic of the sureLDA algorithm.

### Assembling Informative Features

The main ICD and NLP surrogates are counts of the corresponding ICD code(s) and NLP-curated mentions of the phenotype in a patient’s chart. For the *k*th phenotype, we denote these surrogates as *ICD* _*k*_ and *NLP* _*k*_ respectively. Let 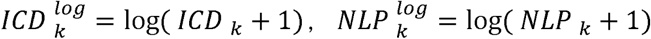, and 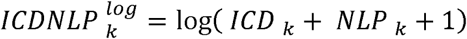. These two key features can be mapped automatically as in Liao et al. [21] using existing knowledge sources including the PheWAS catalogue [12] and the Unified Medical Language System (UMLS). Additional candidate features – including counts of other ICD codes, NLP features, drug prescriptions, lab tests, and procedure codes – can be identified automatically without GLabels via existing methods such as the SAFE method. [23] Since sureLDA has the capacity to select useful features for each phenotype, it is preferable to be inclusive rather than aiming for specificity when assembling these additional features.

### Initializing Prior Probabilities Using PheNorm

We obtain prior probabilities for each of the *K* target phenotypes, denoted by ***π*** = (*π*_1_, …, *π*_*K*_)’, using a slightly modified PheNorm algorithm. In brief, standard PheNorm estimates *π*_*k*_ by (i) normalizing 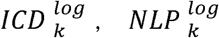 and 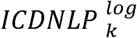 against *H*^*log*^ = log(*H*) via gaussian mixture regression; and (ii) further de-noising these normalized surrogate features using **X**^*log*^ via dropout regression. [20] We use ensemble clustering to estimate *π*_*k*_ by fitting two-class Gaussian mixture models to each of the normalized surrogate features and taking the mean of the predicted probabilities from the three models. For most diseases, patients with *ICD* _*k*_ = 0 - here described as filter negative - very rarely have the disease, so we set *π*_*k*_ = 0 for these patients. The one exception in our study is obesity, which has an unusually insensitive ICD code and for which we define filter negative as *ICD* _*k*_ + *NLP* _*k*_ = 0.

### Fitting Guided LDA

#### Model Overview

Latent Dirichlet Allocation (LDA) is a fully specified topic model that models *D* documents (here patients) as mixtures of *K\** topics (here phenotypes) where each topic is defined as a distribution over a vocabulary of *p* words (here EHR features); for simplicity, we will henceforth discuss the model in terms of patients, phenotypes, and features. To model *K* target phenotypes, we set *K* = K + K*_0_ topics where the first *K* topics are assigned sequentially to target phenotypes 1,.. .,*K* and the remaining *K*_0_ are ‘agnostic’ to model additional structure (i.e non-target phenotypes) in the data. The generative model of standard LDA iterates over three key steps: (i) draw a phenotype mixture ***θ***_*d*_ = (*θ*_1,*d*_,…, *θ*_*K*,d*_)’ for patient d from a Dirichlet distribution with *K\**-dimensional hyperparameter ***α*** = (*α*_1_, …, *α*_*K\**_)’, (ii) draw a feature mixture ***ϕ***_*k*_ = (*ϕ*_*1,k*_,*…, ϕ*_*p,k*_)’ for phenotype topic *k* from a second Dirichlet with *p*-dimensional hyperparameter ***β***, and (iii) for the jth feature, iteratively sample feature-to-phenotype assignment counts ***Z***_*dj*_ = (*Z*_*dj*,1_, …, *Z*_*dj,K\**_)’ from Multinomial (*X*_*dj*_, ***P***_*dj*_) distributions with probability vector ***P***_*dj*_ = (*P*_*dj*,1_,*…, P*_*dj,K\**_)’ calculated based on the current parameter values, where 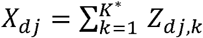 [26] Our guided LDA algorithm resembles the labeled LDA algorithm [29] except whereas labeled LDA requires GLabels to guide topic formation, here we guide topics by setting the Dirichlet hyperparameter for the first *K* phenotype topic distributions to the initial PheNorm probabilities ***π***.

#### Feature Weighting

A major shortcoming of classical LDA is that it weights all terms equally and thus loses precision in the presence of frequent yet uninformative terms called ‘stop words’. [26] To discount these uninformative features, researchers have experimented with term weights including term frequency (TF), inverse document frequency (IDF), the product TF×IDF, and pointwise mutual information. [31] However, these weighting schemes are (1) not reflective of the actual informativeness of a term for a topic, and (2) invariable for a term across topics. In our real EHR data applications, we weight features by the coefficients from the dropout regression step of 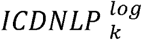 against **X**^*log*^ in PheNorm. [20] Features with a negative dropout regression coefficient for a phenotype are assigned a weight of 0 for the corresponding topic. We henceforth denote the weight of feature *j* for topic *k* as *ω*_*j,k*_

#### Implementation and Inference

We implemented and trained guided LDA using collapsed Gibbs sampling. [32] Collapsed Gibbs sampling is well-suited to sureLDA as we initialize feature-phenotype assignments at their expectations under the PheNorm prior, thus nudging the Gibbs sampler toward a (likely local) optimum within the “vicinity” of the PheNorm solution. Moreover, initialized in this way the Gibbs sampler need only be run once, whereas Variational Bayes would necessitate multiple runs to guarantee finding a reasonably good solution. Our modification of LDA does not substantively affect its Gibbs conditionals, allowing for efficient implementation of this well-trodden procedure. In collapsed Gibbs sampling **Θ** = (***θ***_1_,. .., ***θ***_*D*_) and ***Φ*** = (***ϕ***_1_,. .., ***ϕ***_*K\**_) are marginalized out, so we directly sample the posterior feature-to-disease assignment counts *Z*_*dj*_ without first sampling **Θ** and **Φ**. The algorithm iteratively updates the weighted count of total features assigned to disease topic *k* for patient *d*, 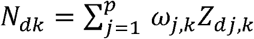, and the weighted count of feature *j* assigned to disease topic *k*, 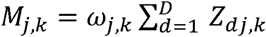, by generating ***Z***_*dj*_ *∼ Multinomial* (*X*_*dj*_, ***P***_*dj*_) where

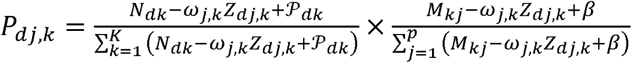

See Algorithm 1 in the Supplementary Materials for details of the algorithm. For the choice of *K\**, we found that including additional ‘agnostic’ topics with uninformative Dirichlet priors (𝒫_*dk*_ = 1) improved performance up to *K*_0_ = 15 additional topics, beyond which performance remained stable up to 100 nadditional topics. We thus set *K\** to *K* + 20.

### Obtaining Final Probabilities Using Ensemble Clustering

When guided LDA converges, we obtain the mean weighted count of total features assigned to topic *k* for subject *d, N*_*dk*_, as the sureLDA score. While **N**_*k*_ = (*N*_*d,k*_,*…, N*_*D,k*_)’ is predictive of **Y**_*k*_, it is subject to noise from healthcare utilization *H*. We therefore obtain our final sureLDA phenotype probability predictions via clustering ensemble normalized by *H*. Specifically, for each *k* we perform a clustering of 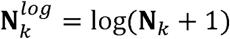 by fitting a gaussian mixture regression model adjusting for *H*^*log*^, and then average this clustering probability with the three PheNorm clustering probabilities based on *H*^*log*^-normalized 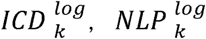, and 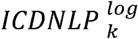.

### Data and Metrics for Evaluation

We evaluated the performance of sureLDA using both simulated datasets and real-world EHR data from the Partners HealthCare Biobank. [33]

#### Simulation Study

In the simulation study, we considered a 5400-dimensional feature space over *K* = 27 disease phenotypes (200 features per disease including 2 surrogate features each) in addition to *H* for *N* = 100,000 patients. To test the robustness of our method’s performance to different phenotype characteristics, we simulated 27 phenotypes enumerating combinations of low, medium and high levels in 1) prevalence, 2) surrogate feature informativeness, and 3) non-surrogate feature informativeness. Mixture models were used to generate **X**, *H* and **Y**. Details of the simulation generative models are given in the Supplementary Materials. The feature weights *ω*_*j,k*_ were uniformly set to 1 in the simulations since no prior knowledge was used in data generation. Results are summarized based on 100 independent and identically distributed simulated datasets.

#### Real EHR Data Analysis

The Partners Healthcare Biobank consists of both codified data (i.e. ICD codes) and free text from clinical notes. We considered 10 target phenotypes - asthma, breast cancer (BrCa), chronic obstructive pulmonary disease (COPD), depression, epilepsy, hypertension (HTN), schizophrenia (SCZ), ischemic stroke (Stroke), type I diabetes mellitus (T1DM), and obesity - to validate our algorithm’s predictive accuracy. These target diseases cover a broad range of acuity, prevalence, and diagnostic ambiguity to test our method’s versatility. We included data from *N* = 38,023 patients observed mostly between 1990 and 2015, when the labels were curated. Moreover, we obtained a total of *p* = 2125 features consisting of age, body mass index (BMI), 110 ICD codes, 225 NLP features counting relevant concepts in clinical notes, all CPT codes grouped into 264 categories according to the Clinical Classifications Software, [34] and 1534 RxNorm drug codes available in the data. NLP features were selected by the SAFE method, while no selection was performed for CPT groups and drug codes. ICD codes were assembled by domain experts in prior studies. Continuous variables such as BMI were rounded to the nearest integer and treated as ordinal in the LDA step. Gold standard labels were manually curated via chart review for 585 patients.

#### Benchmark Methods for Comparison

For each dataset, in addition to fitting sureLDA we considered multiple benchmark approaches: (i) classical LDA automatically assigning latent topics to phenotypes using Spearman’s rank correlation (SRC) with the ICD surrogates, (ii) UPhenome again using SRC with the ICD surrogates to assign topics to phenotypes (only with the Biobank data – we could not apply UPhenome in our simulations as generated features were not assumed to have different datatypes), (iii) GPhenome using *ICD* _*k*_ + *NLP* _*k*_ as the Dirichlet prior for phenotype topic *k*, (iv) the XPress algorithm using *I*(*ICD* _*k*_ ≥ 1) as the proxy label for *Y*_*k*_, (v) PheNorm, and (vi) MAP. We also tested two supervised models: (i) LASSO-penalized logistic regression and (ii) random forest, each with *n*_*t*_ training labels where we let *n*_*t*_ vary from 100-300 to assess how sureLDA compares to supervised phenotyping methods. In light of Rajkomar et al.’s recent result that LASSO logistic regression rivals deep learning methods on a set of high-dimensional EHR-based prediction tasks, and given the limited number of GLabels, we did not include more complex supervised methods. [35] Hyperparameters for LASSO and random forest were optimized using 10-fold cross-validation. 2- and 5-fold cross-validation as well as random subsampling cross-validation yielded statistically equivalent optimized hyperparameters and predictive accuracy for all phenotypes in the Partners EHR dataset (data not shown).

#### Evaluation Metrics

We used three evaluation metrics to quantify the predictive performance of sureLDA and its comparators: (i) area under the receiver operator curve (AUC), (ii) *F* score, and (iii) label rank loss. [36] For *F* score, we chose the cutoff value to achieve a specificity of 95% among filter-positive samples. AUC and F score reflect the sensitivity/specificity and precision/recall of predictions respectively. On the other hand, label rank loss computes the average number of probabilistic label pairs for a patient that are incorrectly ordered weighted by the inverse of the number of ordered pairs of false and true labels, reflecting the degree to which the relative disease probabilities for a given patient align with that patient’s true overall phenotype. Thus, whereas the first two evaluation metrics measure the quality of predictions for a phenotype, the third measures accuracy on the patient level and thereby reflects a model’s capacity to jointly predict multiple phenotypes. Standard errors for all measurements were obtained by bootstrapping with 100 bootstrap samples. To evaluate methods’ robustness to diverse phenotypes, we considered the standard deviation of Δ = *AUC*_*max*_ − *AUC*_*method*_ across Partners Biobank phenotypes, where for each phenotype *AUC*_*max*_ denotes the maximum AUC achieved across label-free phenotyping methods, and *AUC*_*method*_ denotes the AUC of the specific method. Finally, for the Partners EHR data, we qualitatively assessed whether the phenotype topics inferred in the guided LDA step make clinical sense by generating feature clouds using the weighted counts of features assigned to each phenotype topic, {*M*_*j,k*_}.

## RESULTS

### Results on Simulated Datasets

Figure 2 shows mean accuracies across 27 simulated phenotypes with variable generative parameters. Figure 3 shows AUCs and *F* scores as a function of (a) prevalence, (b) surrogate feature informativeness, and (c) non-surrogate feature informativeness. sureLDA, classical LDA, and GPhenome’s accuracy (per both AUC and *F* score) improved significantly with the informativeness of both surrogate and non-surrogate features. MAP improved only with surrogate informativeness as expected. PheNorm and Xpress improved markedly with surrogate informativeness and marginally with non-surrogate informativeness. Finally, increasing phenotype prevalence improved □ scores but not AUCs (which as a metric is prevalence-agnostic) for all methods except XPress.

**Figure 2:**
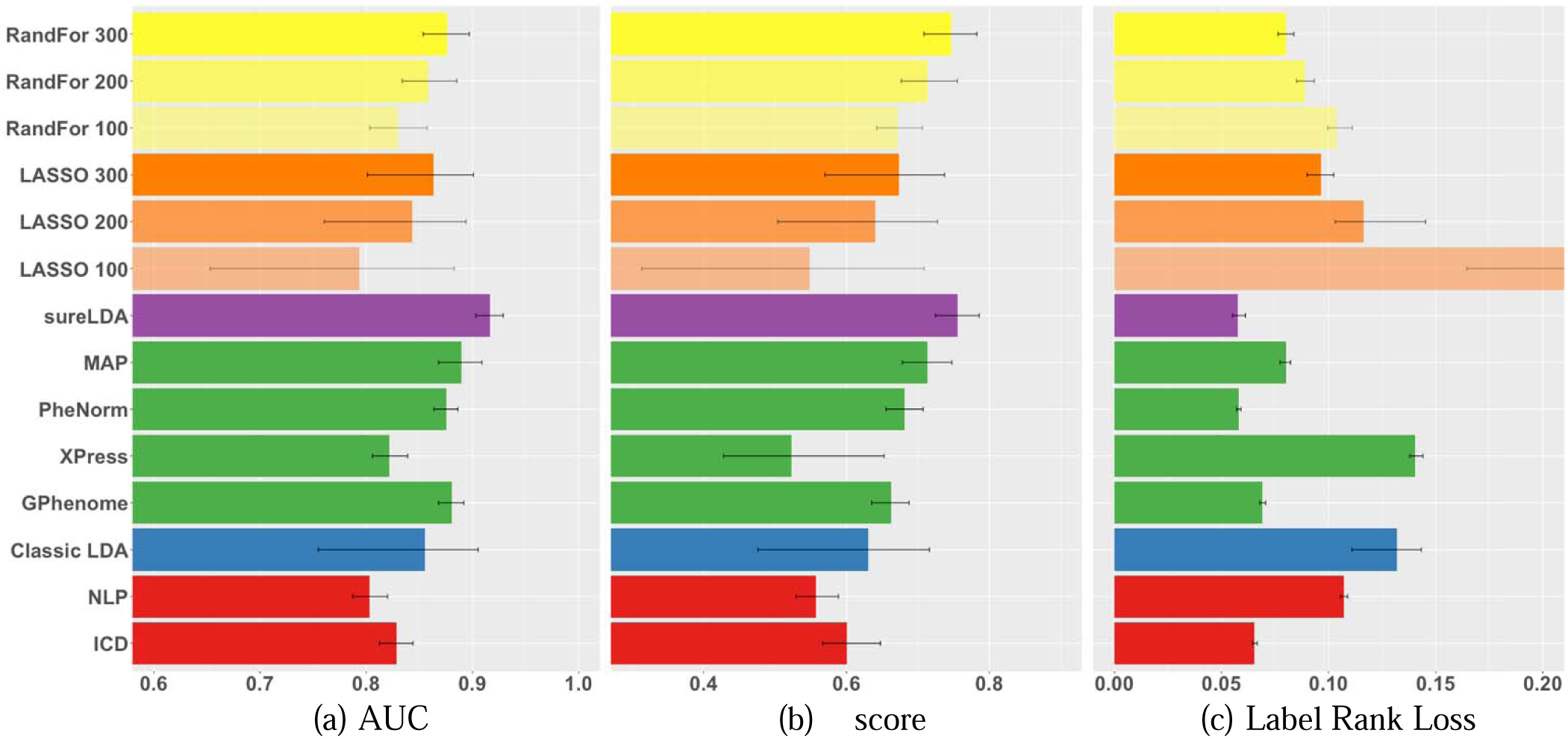
Mean (a) AUCs, (b) *F* scores, and (c) label rank losses of phenotype predictions on simulated datasets comparing raw ICD and NLP surrogates (red), fully unsupervised phenotyping methods (blue), alternative weakly supervised methods (green), sureLDA (purple), and supervised phenotyping using LASSO regularized logistic regression (LASSO, orange) and random forest (RandFor, yellow) with 100-300 true labels. Error bars reflect empiric bootstrapped 95% confidence intervals.

**Figure 3:**
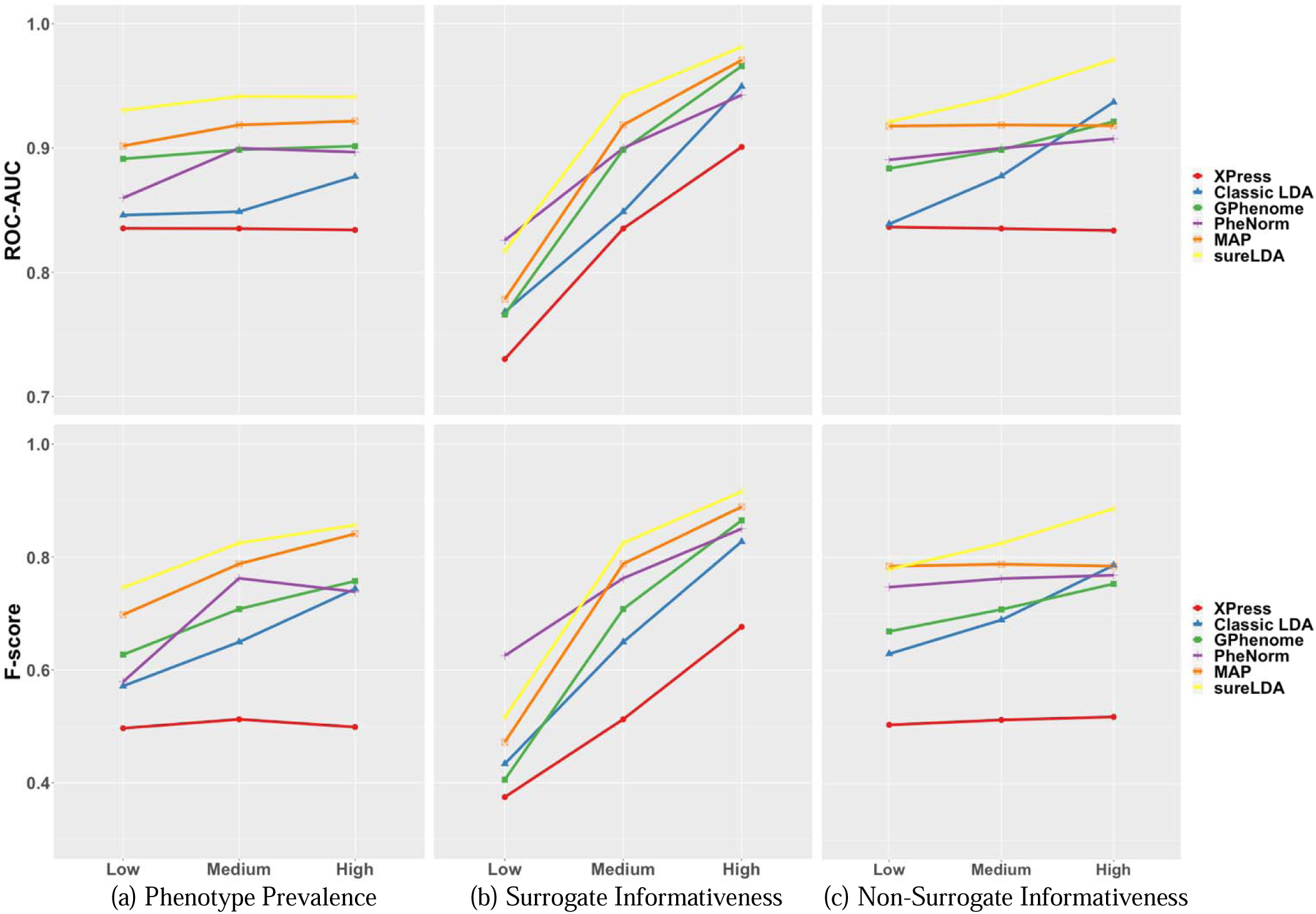
Mean AUCs (top panel) and *F* scores (bottom panel) of Xpress (red), classic LDA (blue), GPhenome (green), PheNorm (purple), MAP (orange), and sureLDA (yellow) on simulated datasets varying (a) phenotype prevalence, (b) surrogate feature informativeness, and (c) non-surrogate feature informativeness. For each plot, the two variables not being varied are held at their ‘Medium’ levels.

### Results on Partners EHR Data

Figure 4 shows mean accuracies across 10 diverse diseases in real-world EHR data. sureLDA exhibited statistically significant improvements in mean AUC relative to other label-free alternatives. For example, compared to MAP, PheNorm, Xpress, and GPhenome, sureLDA improved mean AUCs by 0.021 (95% CI: [0.004, 0.038]), 0.033 (95% CI: [0.008, 0.052]), 0.131 (95% CI: [0.100, 0.170]), and 0.054 (95% CI: [0.028,0.084]) respectively. Compared to supervised algorithms, sureLDA achieved statistically significant improvements in mean AUC as well as improvements (though not significant) in mean *F* score relative to both LASSO and Random Forest with 300 GLabels. sureLDA also exhibited relatively low standard errors in all accuracy metrics. As shown in Figure S3, sureLDA’s deviation from the highest AUC achieved per disease had the lowest standard deviation (0.029), with only PheNorm (0.044) and GPhenome (0.046) achieving comparable robustness by this metric. These results were generally consistent with our simulation study.

**Figure 4:**
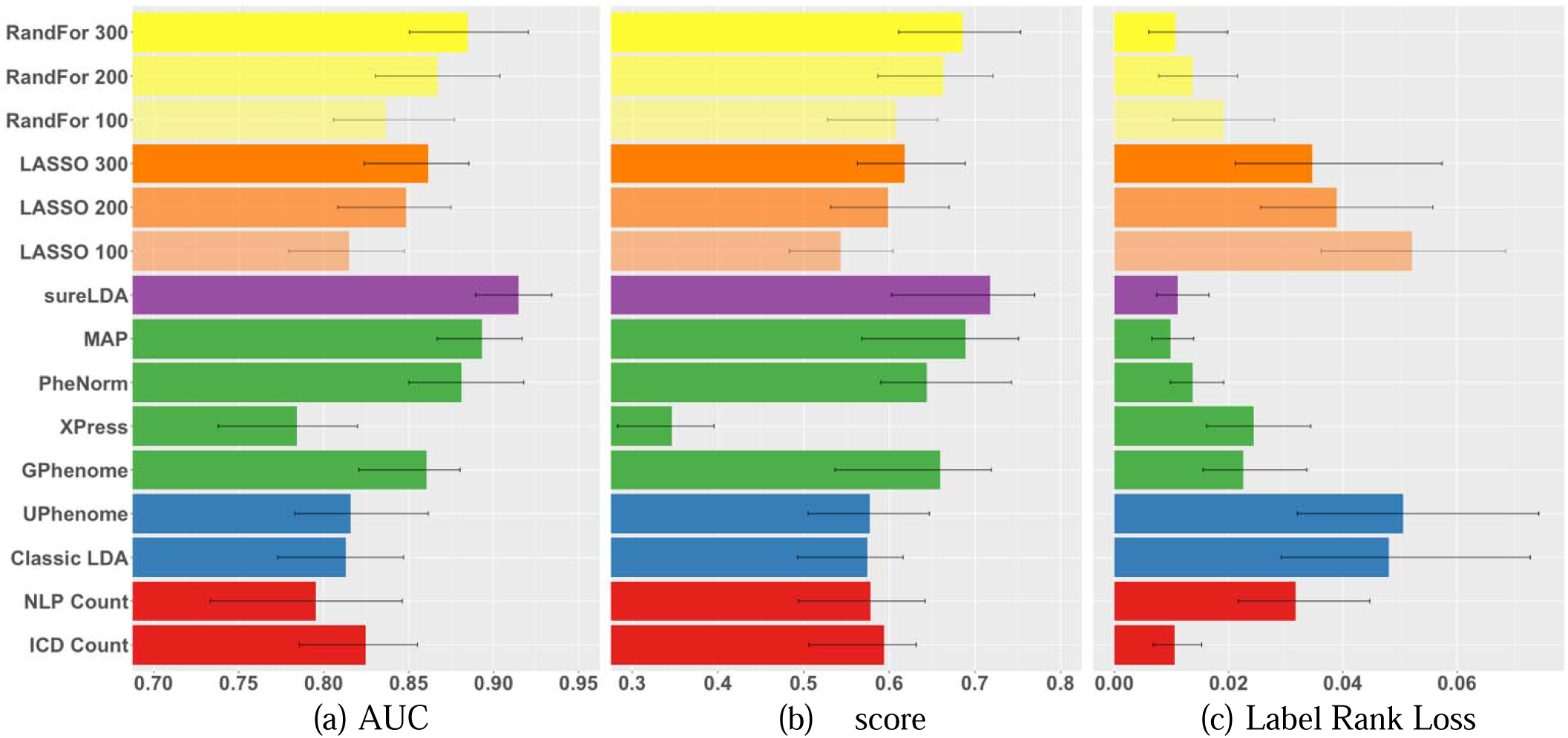
Mean (a) AUCs, (b) *F* scores, and (c) label rank losses of phenotype predictions on real diseases from the Partners EHR Biobank comparing raw ICD and NLP surrogates (red), fully unsupervised phenotyping methods (blue), alternative weakly supervised methods (green), sureLDA (purple), and supervised phenotyping using LASSO regularized logistic regression (LASSO, orange) and random forest (RandFor, yellow) with 100-300 true labels. Error bars reflect empiric bootstrapped 95% confidence intervals.

Per the label rank loss metric, which reflects the accuracy of relative phenotype predictions on the patient level, sureLDA achieved statistically equal loss to MAP, PheNorm, ICD counts, and Random Forest with 300 labels. GPhenome, UPhenome, and classic LDA’s losses are confounded by the fact that they output topic-feature counts – which vary in scale between phenotypes – rather than probabilities. These results depart from our simulation, in which sureLDA achieved the lowest loss statistically equivalent only to PheNorm. This disparity may be attributable to the fact that our simulation has 27 phenotypes whereas our real-world EHR example has only 10, which may not be sufficient to observe the benefit of jointly modeling phenotypes over predicting each marginally. Nevertheless, sureLDA’s low rank loss in our simulation study, combined with the qualitative observation that sureLDA consistently achieves higher mean AUCs and *F* scores the more phenotypes we include (data not shown), suggests that sureLDA benefits from jointly modeling phenotypes.

sureLDA performed statististically equivalently to or better than MAP and PheNorm for all phenotypes, suggesting that its anchoring to the PheNorm prior ensures reliable baseline accuracy. Consistent with our simulation results, sureLDA generally performed best relative to alternative label-free methods for phenotypes with informative non-surrogate features and at least one noisy surrogate, such as COPD, epilepsy, and ischemic stroke (Figures S1-S2, Table S1). However, sureLDA along with MAP and PheNorm were significantly outperformed by GPhenome for obesity – which has unusually insensitive ICD and NLP surrogates. This suggests that sureLDA’s tethering to PheNorm may occasionally hinder it from fully exploiting information distributed across non-surrogate features.

We present in Figures 5 and S5 feature clouds demonstrating the 20 features sureLDA associates most closely with each target phenotype “topic.” Almost all of these features have an intuitive association with their corresponding phenotype. For instance, highly-weighted features for the epilepsy topic include the ICD code for epilepsy, anticonvulsive drugs including phenytoin, lamotrigine, and carbamazepine, the diagnostic test electroencephalogram (EEG), and various NLP terms including “seizures” and “aura” – a multitype set of features clearly reflecting a diagnosis of epilepsy (Figure 5a). These results underline sureLDA’s robust feature selection properties in the high-dimensional data setting,

**Figure 5:**
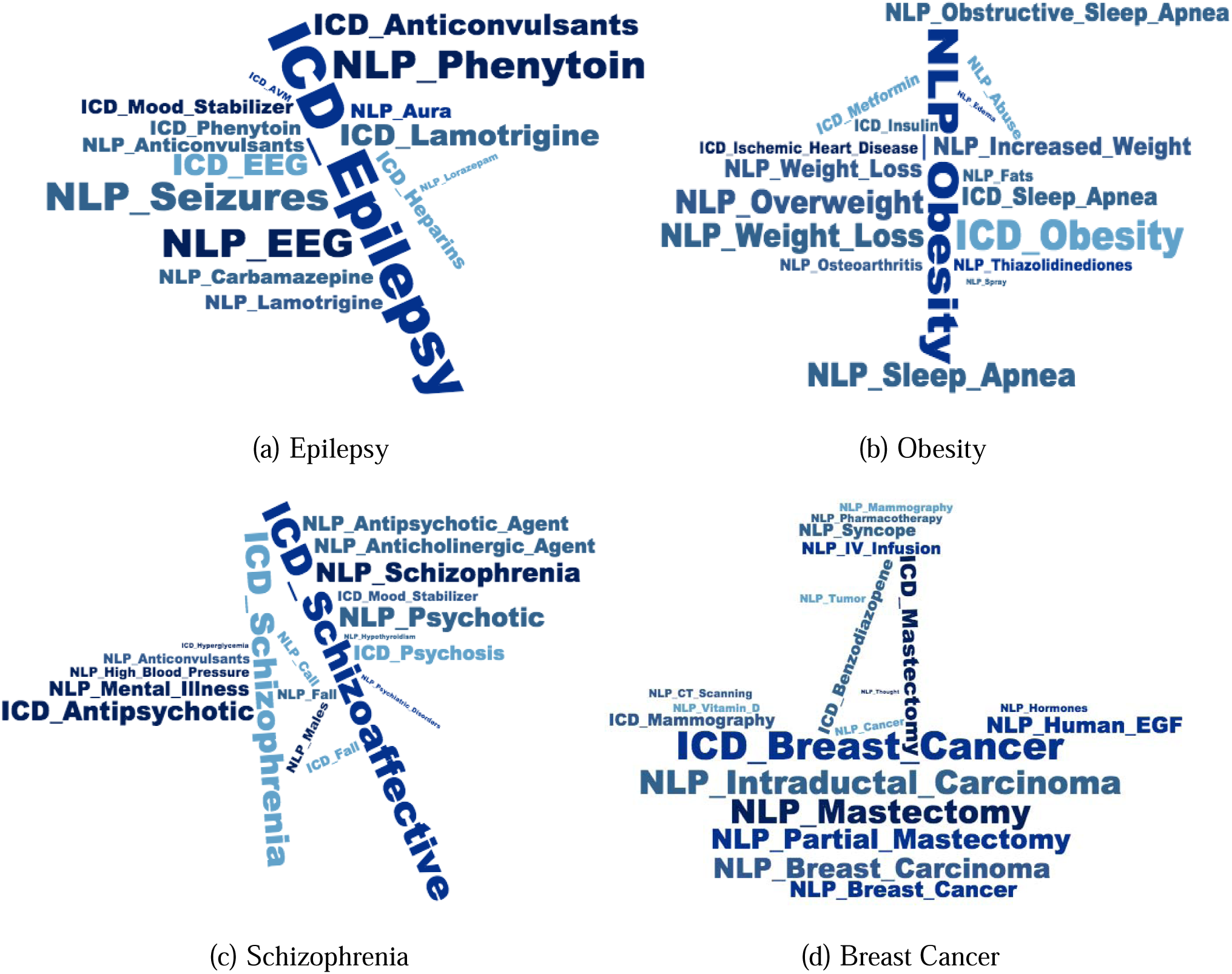
Feature clouds derived from *M*_*j,k*_ for four representative disease phenotypes: epilepsy, obesity, schizophrenia, and breast cancer. NLP terms have the prefix ‘NLP’ and codified data the prefix ‘ICD’.

## DISCUSSION

Automated, label-free phenotyping methods enable EHR studies for which manually annotating phenotypes or designing new rule-based algorithms are impractical, such as a PheWAS requiring a large number of phenotypes to be defined and labeled *de novo*. To this end, sureLDA enables efficient annotation of accurate, interpretable computational phenotype labels that are robust to phenotype prevalence and feature set properties (Figures 3, S3).

sureLDA combines attractive properties of classical LDA and PheNorm. By leveraging PheNorm to guide topic formation in LDA in a weakly supervised manner, sureLDA focuses LDA’s structure identification prowess towards identifying a well-defined set of phenotypes. It also leverages PheNorm’s dropout regression step to enable a data-driven phenotype-specific feature weighting mechanism that outperforms the literature-standard TF-IDF weighting scheme (Figure S4).

Likewise, sureLDA augments PheNorm by using LDA to better extract information from non-surrogate features. LDA is adept at attributing marginally ambiguous features such as ‘cough’ or ‘headache’ to the most patient-relevant phenotype topic. In this way, it implicitly imposes an Occam’s Razor assumption that favors fewer concurrent phenotypes over more, reflecting the negative association that arises between phenotypes conditional on ambiguous features. Future work is warranted to better assess the benefit of this property in the real-world EHR setting.

LDA also gives sureLDA its aptitude for feature selection. As the word clouds in Figures 5 and S5 demonstrate, the sureLDA ‘scores’ produced by guided LDA reflect consistently meaningful features. Given the noisy and high-dimensional nature of EHR data, sureLDA’s feature selection ability primes it for EHR modeling. Moreover, it enables fast production of interpretable computational phenotypes.

This combination of desirable properties from PheNorm and LDA explains sureLDA’s robustness to a diversity of phenotypes (Figures 3, S3). For phenotypes with highly informative surrogates, the use of PheNorm ensures high baseline accuracy. For phenotypes with noisier surrogates and informative non-surrogate features, the LDA step allows for significant improvement over PheNorm. This robustness makes sureLDA well-suited to simultaneously annotating many diverse phenotypes.

sureLDA could also be used to identify subphenotypes similarly to Li et al. [37] Once a phenotype (i.e. heart failure) cohort is identified (potentially via sureLDA), one could employ sureLDA on these patients to classify subphenotypes. However, since sureLDA uses PheNorm to guide topics, one would need to prespecify a few known subphenotypes (i.e. systolic heart failure with renal complications) and use their relevant ICD and NLP features to guide the subphenotype topics. Additionally, sureLDA’s ‘agnostic’ topics could be used to discover unknown (sub)phenotypes, though its advantage over standard LDA for this task is unclear. Phenotype discovery is also limited by what information is documented in the EHR, which inherently reflects known phenotypes. While sureLDA eliminates reliance on expert review for annotation, it does require gold-standard labels for evaluation of its own performance. Continued work is warranted to estimate performance parameters such as AUC and F-score in an unsupervised fashion and thereby make both implementation and evaluation of weakly supervised phenotyping methods like sureLDA fully label-independent.

## CONCLUSION

In this paper we introduce sureLDA, a high-throughput automated EHR phenotyping method that combines PheNorm with LDA to jointly annotate multiple phenotypes without using GLabels. Our method produces accurate, interpretable labels for a broad range of phenotypes. sureLDA exhibits particular improvement over existing label-free phenotyping methods for phenotypes without accurate surrogate features. Given these qualities, sureLDA promises to enable more powerful use of EHR data for highly multi-phenotype applications such as PheWAS.

## Contributorship

All authors made substantial contributions to: conception and design; acquisition, analysis and interpretation of data; drafting the article or revising it critically for important intellectual content; and final approval of the version to be published.

