## Supplemental Materials for "sureLDA: A Multi-Disease Automated Phenotyping Method for the Electronic Health Record"

**Supplementary Materials**

### 1. Guided LDA Algorithm

The guided LDA procedure is summarized in Algorithm 1. In practice, we ran R=4 randomly-initialized Gibbs sampling chains for $N_{R}$=200 iterations apiece, discarded the first 50 iterations of each chain as burn-in, and estimated $\mathbb{N=[}N_{dk}]_{D\times K^{*}}$ and $\mathbb{M=[}M_{j,k}]_{K^{*}\times J}$ as the mean of the 600 remaining samples for each respective parameter. In keeping with common practices in inferential sampling, we assessed convergence of our estimates as having an inter-chain potential scale reduction $<1.05$.[1] We selected a burn-in by qualitatively analyzing individual $N_{dk}$ and $M_{j,k}$ samplers, finding consistent convergence to stationary distributions within 25 iterations. To ensure that variance from the Gibbs sampling procedure did not unduly influence our reported performance estimates, we ran 100 instances of the above procedure and empirically estimated the Gibbs-sampling-attributable standard error in the predictive AUC, *F* score, and rank loss estimates. We found these errors to be reliably in the $[0.001,0.002]$ range - functionally insignificant compared to other sources of random error in the overall sureLDA procedure. We heuristically set the Dirichlet hyperparameters $\alpha$ and $\beta$ to 1 to reflect their prior uninformativeness.

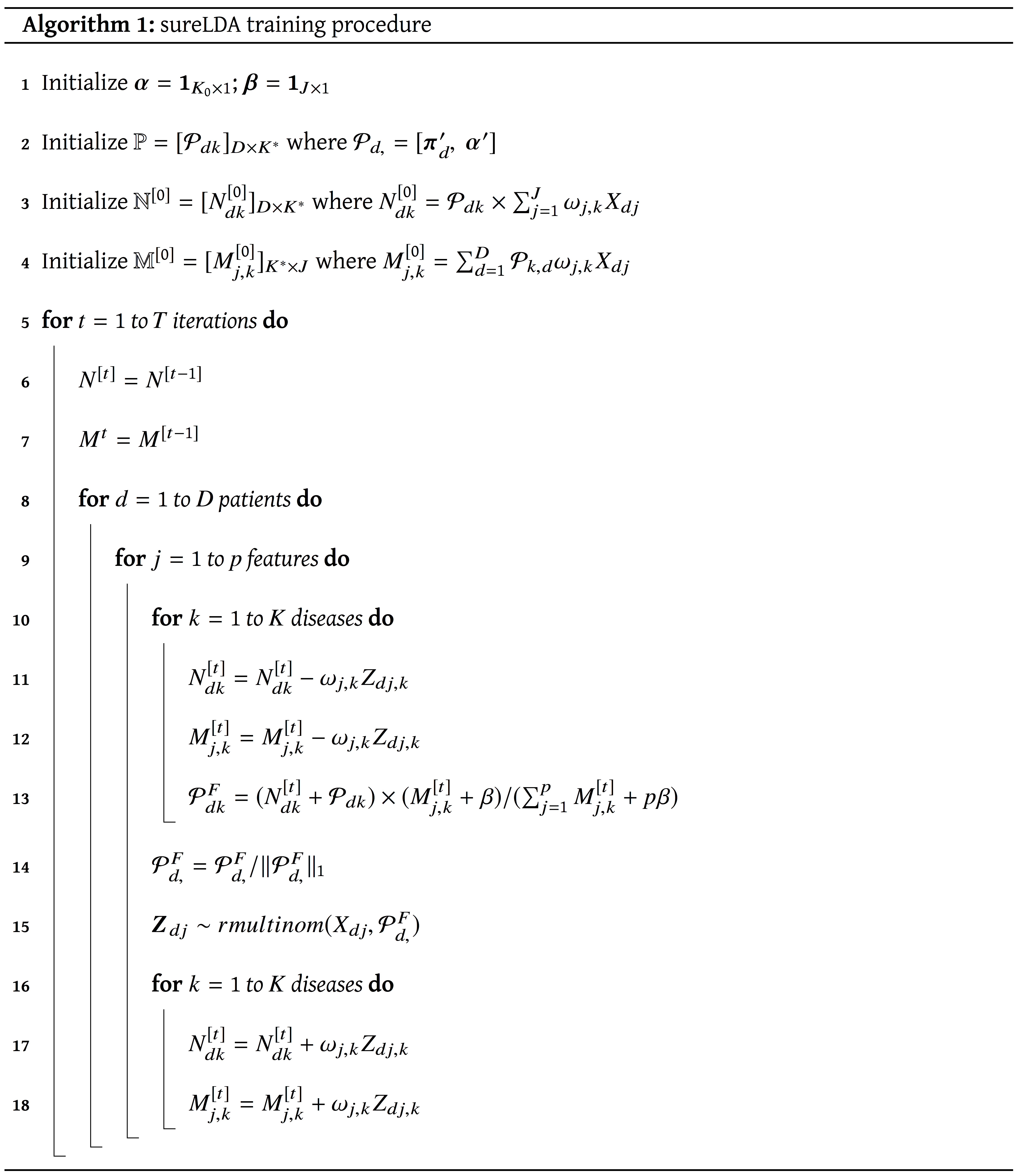

### 2. Details of Simulation Set-up

We first generate $\mathbf{Y}=(Y_{1},...,Y_{10})'$ with $Y_{k}\in\{+,-\}$ from correlated multivariate binary random variables by thresholding multivariate normal random vectors with mean $(.4,.2,.1,.1,.08,.07,.06,.04,.02,.01)'$ and variance 0.25. We let the correlation of the underlying multivariate normal be $\rho=0.5$ for diseases 1 and 2, and $\rho=0$ for all other pairs. We let $p=5000$, the feature vector $\mathbf{X}=(H,\mathbf{x}_{1'},...,\mathbf{x}_{10}')'$, and generated $\{\mathbf{x}_{k}=( ICD{}_{k}, NLP{}_{k},x_{k,1},...,x_{k,498})',k=1,...,K\}$ as follows. We first generate $H\sim Poisson (\lambda=2)$, and then $\{ ICD{}_{k}^{*}, NLP{}_{k}^{*},x_{k,ȷ},k=1,...,10,ȷ=1,...,498\}$ from

$( ICD{}_{k}^{*}+1)(H+1)^{-0.30}|\mathbf{Y}\sim Gamma (\beta_{ICD_{k}}^{Y_{k}},1),$

$( NLP{}_{k}^{*}+1)(H+1)^{-0.25}|\mathbf{Y}\sim Gamma (\beta_{NLP_{k}}^{Y_{k}},1),$

$x_{k,ȷ}^{*}(H+1)^{-0.20}|\mathbf{Y}\sim Gamma (b_{k,ȷ}^{Y_{k}},1), k=1,..,10,ȷ=1,...,498,$

where for $k=1,...,10$ and $ȷ=1,...,498$

$\beta_{ICD_{k}}^{+}=5.5^{I(k\leq5)}4.5^{I(k\geq6)}, \beta_{NLP_{k}}^{+}=2.1, b_{k,j}^{+}=1.25$

$\beta_{ICD_{k}}^{-}=0.71^{I(k\leq5)}0.51^{I(k\geq6)}, \beta_{NLP_{k}}^{-}=0.81, b_{k,j}^{-}=0.9^{I(k\geq6)}.$

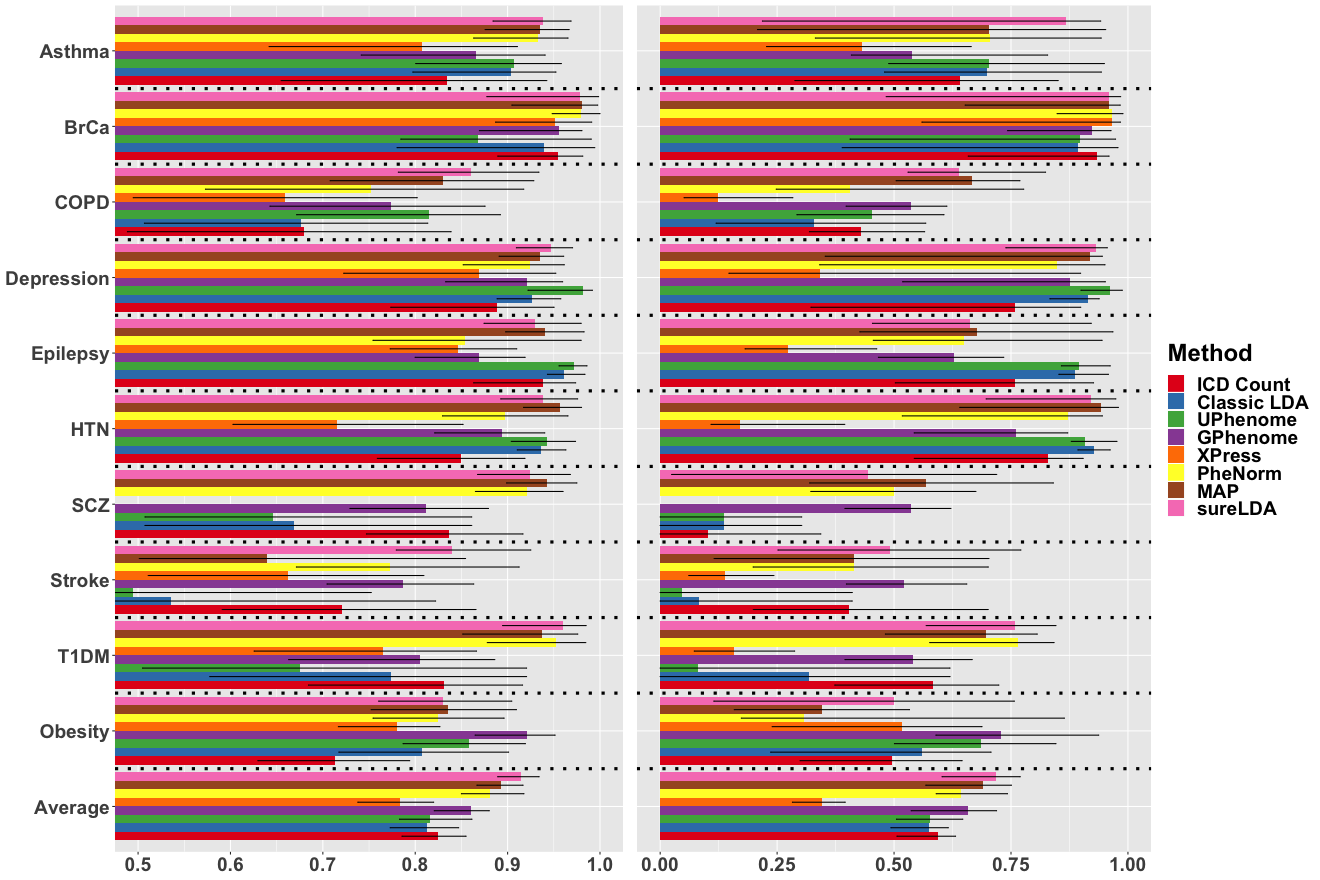

(a) AUC	(b)$F$ score

**Figure S1**: AUCs and $F$ scores of phenotype predictions on 10 disease phenotypes in the Partners EHR Biobank by sureLDA versus label-free comparators including $ICD{}_{k}$, classic LDA, UPhenome, GPhenome, XPress, PheNorm, and MAP. Error bars reflect empiric bootstrapped $95\%$ confidence intervals.

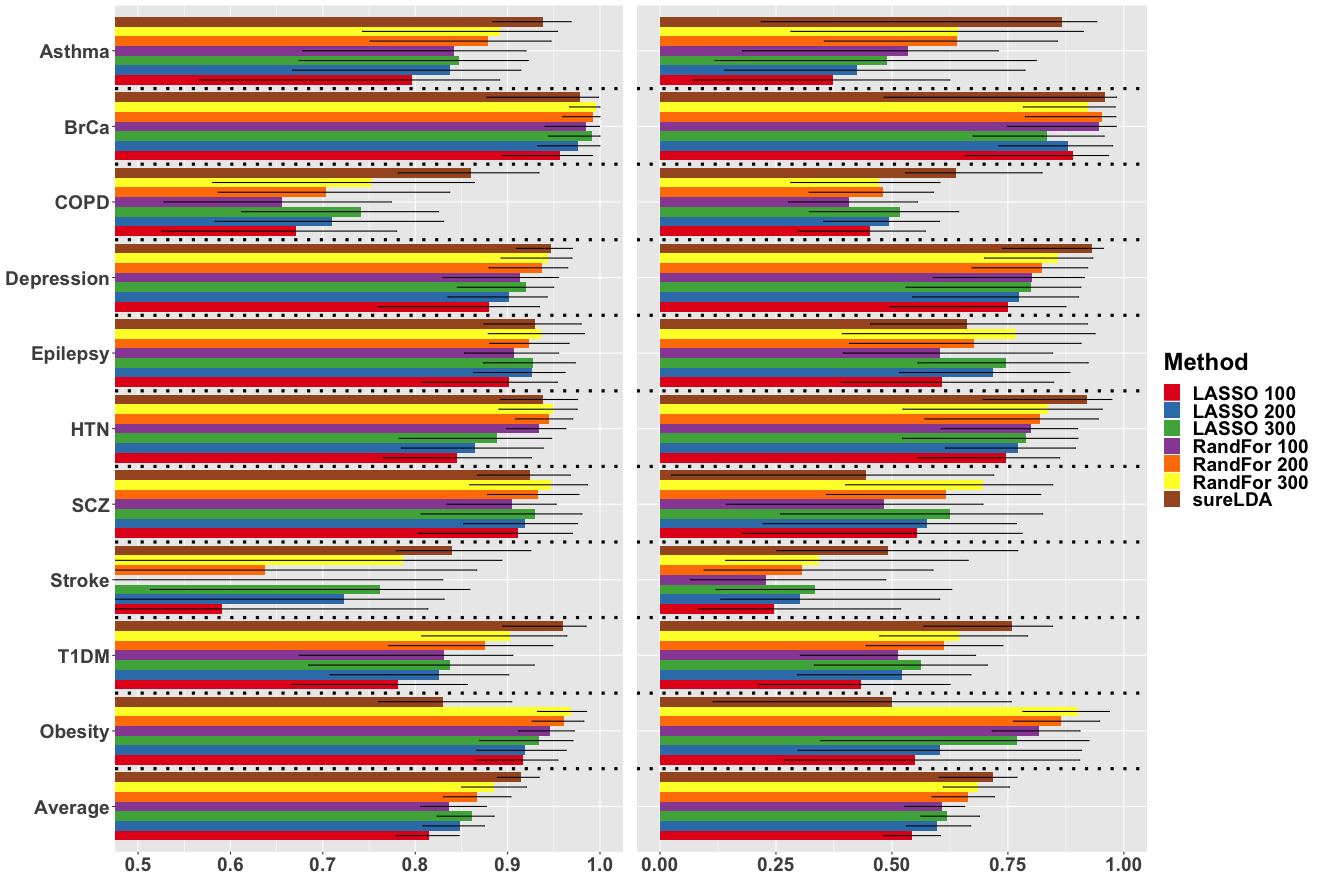

(a) AUC	(b)$F$ score

**Figure S2**: AUCs and $F$ scores of phenotype predictions on 10 disease phenotypes in the Partners EHR Biobank by sureLDA versus LASSO regularized logistic regression (LASSO) and random forest (RandFor) each trained on 100-300 samples. Error bars reflect empiric bootstrapped $95\%$ confidence intervals.

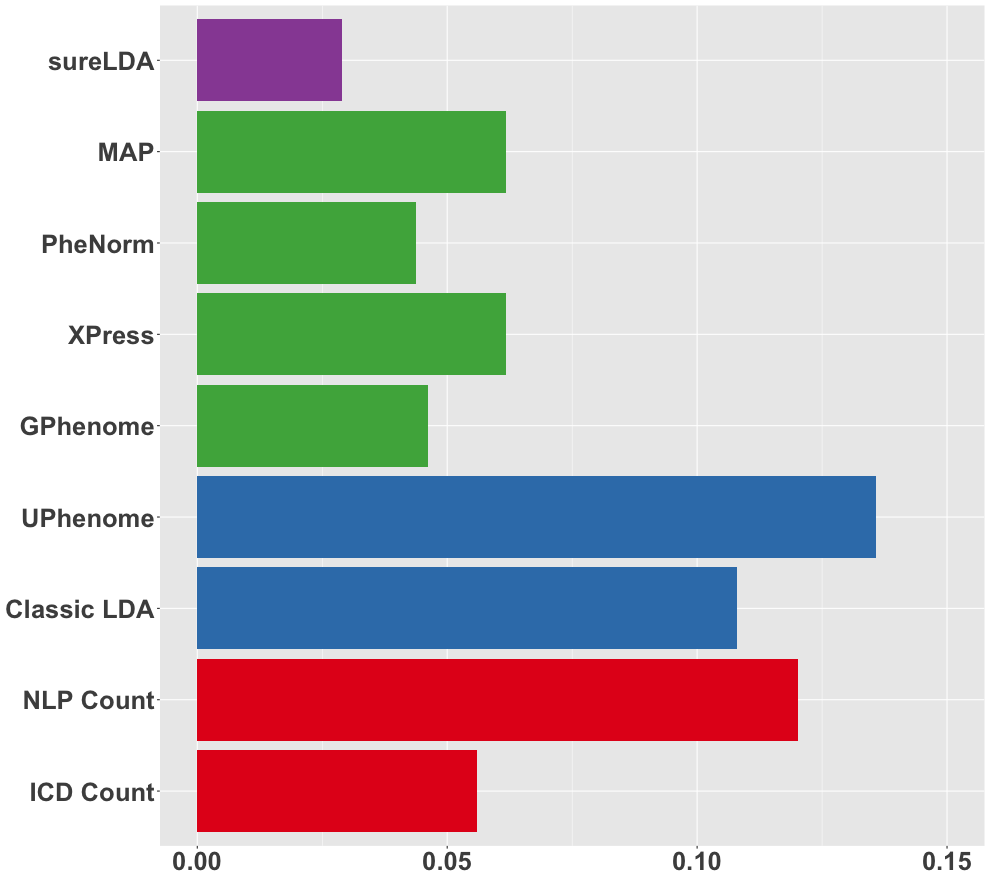

**Figure S3:** Standard deviations of the delta between a method’s AUC for a Partners EHR phenotype and the maximum AUC achieved for that phenotype across label-free phenotyping methods, comparing sureLDA (purple) to raw ICD and NLP surrogates (red), fully unsupervised methods (blue), and alternative weakly supervised methods (green).

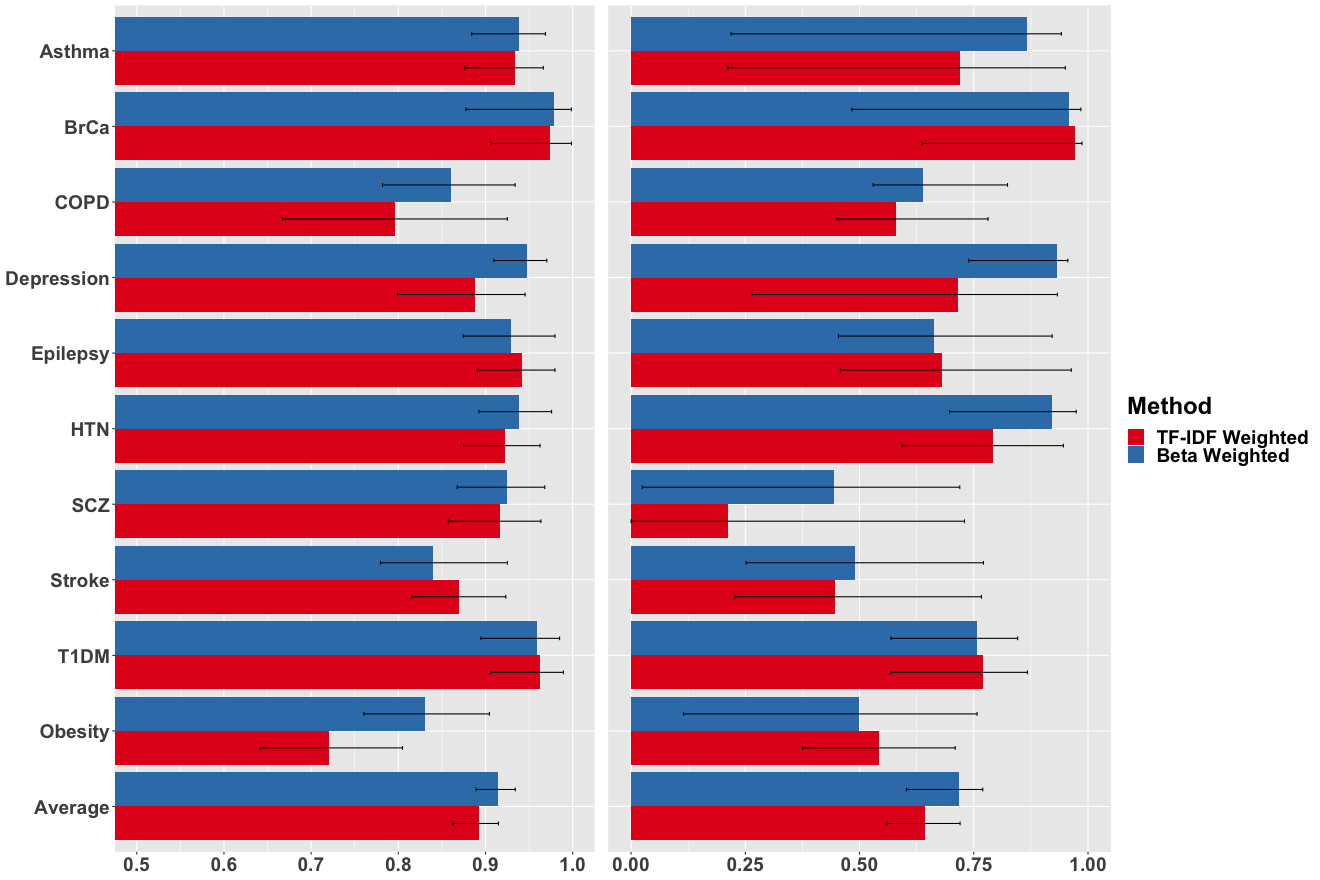

(a) AUC	(b) $F$ score

**Figure S4:** AUCs and *F* scores of sureLDA phenotype predictions for 10 disease phenotypes in the Partners EHR Biobank, weighting terms in the guided LDA step using the unbiased TF-IDF scheme versus the PheNorm beta coefficient scheme introduced in this study. Error bars reflect empiric bootstrapped $95\%$ confidence intervals.

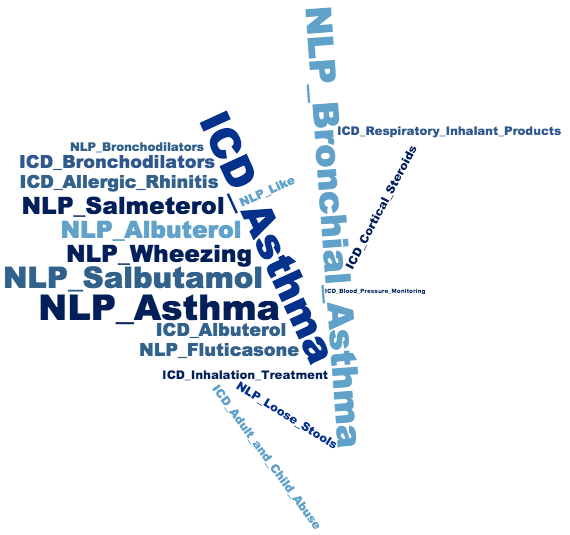

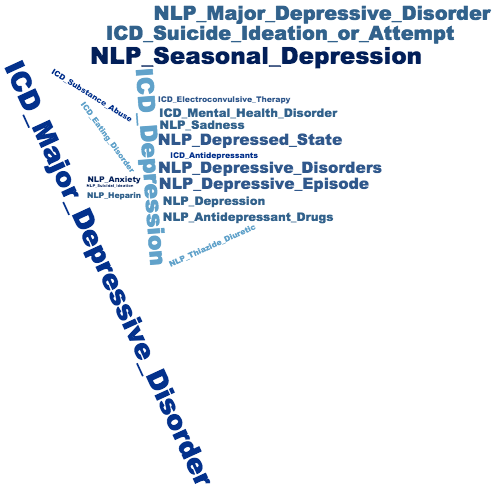

(a) Asthma	(b) Depresssion

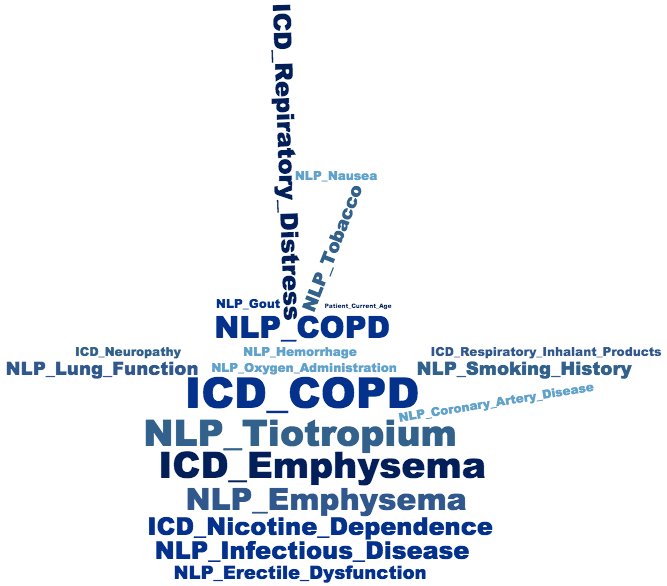

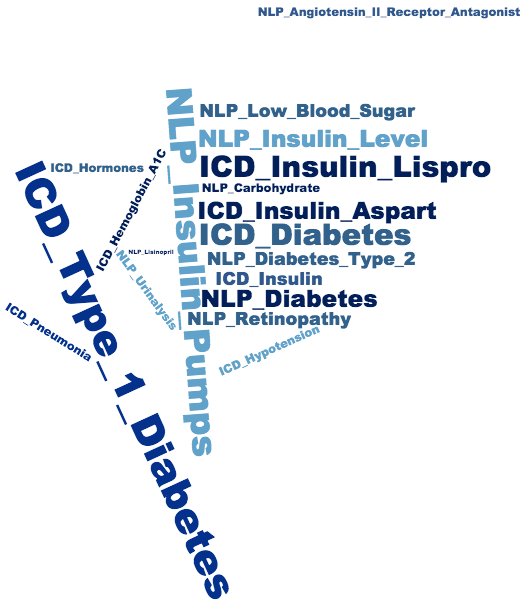

(c) COPD	(d) Type 1 Diabetes

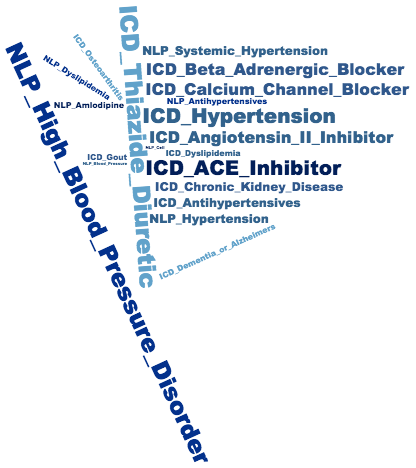

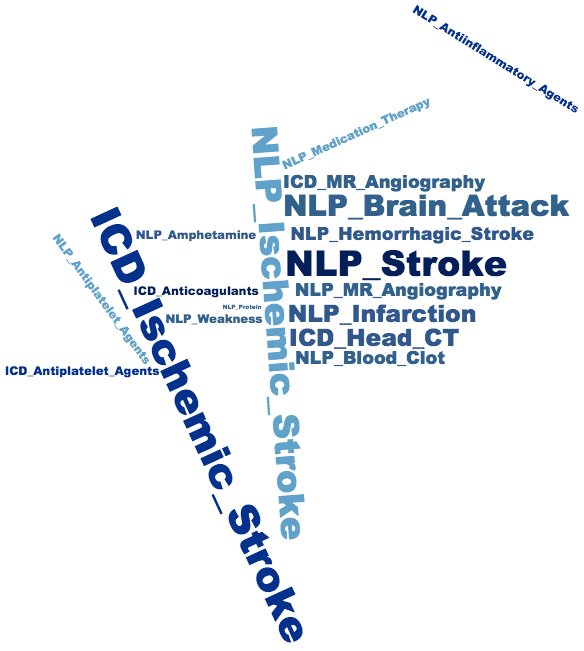

(e) Hypertension	(f) Ischemic Stroke

**Figure S5:** Feature clouds derived from $M_{j,k}$ for (a) asthma, (b) depression, (c) COPD, (d) type 1 diabetes mellitus, (e) hypertension, and (f) ischemic stroke.

| Estimator | Asthma | BrCa | COPD | Depres | Epilep | Hypert | Schizo | Stroke | T1DM | Obesity | Mean | SD(Δ) |
| --- | --- | --- | --- | --- | --- | --- | --- | --- | --- | --- | --- | --- |
| ICD | .834 (.083) | .955 (.024) | .679 (.099) | .889 (.046) | .938 (.029) | .849 (.043) | .836 (.042) | .721 (.075) | .831 (.060) | .713 (.041) | .825 (.018) | .056 |
| NLP | .857 (.060) | .950 (.049) | .794 (.051) | .890 (.047) | .664 (.159) | .843 (.066) | .935 (.025) | .506 (.095) | .677 (.093) | .837 (.034) | .795 (.030) | .120 |
| Classic LDA | .903 (.046) | .939 (.033) | .677 (.094) | .926 (.104) | .961 (.045) | .936 (.065) | .669 (.061) | .536 (.185) | .773 (.084) | .807 (.034) | .813 (.030) | .108 |
| UPhenome | .907 (.047) | .868 (.033) | .815 (.059) | **.982 (.042)** | **.971 (.032)** | .943 (.076) | .646 (.081) | .494 (.161) | .675 (.100) | .858 (.024) | .816 (.027) | .136 |
| GPhenome | .865 (.051) | .956 (.026) | .774 (.059) | .921 (.030) | .869 (.025) | .894 (.029) | .812 (.031) | .787 (.037) | .806 (.052) | **.921 (.019)** | .860 (.014) | .046 |
| XPress | .807 (.085) | .951 (.027) | .660 (.087) | .869 (.064) | .846 (.037) | .715 (.065) | NA | .662 (.078) | .765 (.064) | .780 (.029) | .784 (.021) | .062 |
| PheNorm | .933 (.027) | .980 (.015) | .752 (.105) | .924 (.030) | .854 (.073) | .897 (.035) | .921 (.026) | .773 (.063) | .953 (.029) | .824 (.039) | .881 (.019) | .044 |
| MAP | .935 (.025) | **.981 (.026)** | .830 (.064) | .935 (.019) | .941 (.023) | **.957 (.017)** | **.943 (.021)** | .639 (.104) | .937 (.035) | .835 (.041) | .893 (.013) | .062 |
| sureLDA | **.938 (.023)** | .978 (.031) | **.860 (.047)** | .947 (.016) | .929 (.029) | .938 (.023) | .924 (.027) | **.839 (.040)** | **.960 (.025)** | .830 (.039) | **.914 (.011)** | **.029** |
| LASSO${}_{100}$ | .797 (.082) | .956 (.028) | .671 (.065) | .880 (.043) | .901 (.035) | .846 (.043) | .911 (.039) | .591 (.126) | .781 (.054) | .916 (.025) | .815 (.019) | N/A |
| LASSO${}_{200}$ | .837 (.059) | .977 (.020) | .709 (.063) | .901 (.028) | .927 (.029) | .865 (.041) | .918 (.034) | .723 (.101) | .825 (.053) | .918 (.027) | .848 (.018) | N/A |
| LASSO${}_{300}$ | .847 (.065) | .991 (.017) | .741 (.060) | .920 (.028) | .928 (.030) | .888 (.048) | .929 (.043) | .762 (.097) | .837 (.061) | .934 (.029) | .861 (.017) | N/A |
| RandFor${}_{100}$ | .842 (.063) | .985 (.018) | .656 (.065) | .913 (.034) | .906 (.026) | .935 (.018) | .905 (.033) | .431 (.185) | .831 (.059) | .946 (.016) | .837 (.021) | N/A |
| RandFor${}_{200}$ | .879 (.057) | .993 (.012) | .703 (.068) | .937 (.024) | .923 (.024) | .944 (.018) | .933 (.030) | .638 (.188) | .876 (.046) | .961 (.015) | .867 (.021) | N/A |
| RandFor${}_{300}$ | .892 (.058) | .996 (.010) | .753 (.075) | .944 (.021) | .937 (.027) | .949 (.022) | .947 (.032) | .787 (.176) | .902 (.042) | .969 (.014) | .885 (.019) | N/A |

**Table S1**: Columns 1-11 contain AUCs (with empiric bootstrapped standard errors) of the main ICD and NLP surrogates, XPress, classic LDA, UPhenome, GPhenome, PheNorm, MAP, sureLDA, LASSO regularized logistic regression (LASSO), and random forest (RandFor) with 100-300 training labels respectively, on 10 real diseases from the Partners EHR Biobank. Column 12 contains standard deviations of the delta between a method’s AUC for a phenotype and the maximum AUC achieved for that phenotype across label-free phenotyping methods (see *Evaluation Metrics* subsection of *Methods* for details). The highest AUC (and lowest SD(Δ)) achieved via label-free methods for each disease is boldfaced.
